# Sequential prioritization of expected and unexpected information in visual processing

**DOI:** 10.64898/2026.03.23.713176

**Authors:** Arjen Alink, Janika Becker, Helen Blank

## Abstract

This electroencephalography (EEG) study provides evidence that expectations dynamically shape visually evoked brain responses through a biphasic temporal mechanism: an earlier enhancement of expected image components, followed by a later prioritization of unexpected components. Across six sessions each, six participants were presented with a deterministic sequence of four images with intermixed composite trials that contained both the expected and the unexpected image. To quantify expected and unexpected identity information over time, we applied Linear Discriminant Analysis. The observed temporal dissociation suggests that the brain initially integrates sensory input with prior expectations in a Bayesian manner before shifting to emphasize surprising information, thereby supporting both efficient recognition and adaptive model updating. Furthermore, expectations affected superordinate image processing (animal vs. non-animal), showing the same pattern as identity processing, with earlier enhancement of expected information followed by a prioritization of surprising information.

## Introduction

Several decades ago, a paper by Rao and Ballard^1^ reinvigorated the Helmholtzian idea that perception arises from a synergy between sensations and expectations. Since then, a large cohort of human neuroimaging studies has interpreted contextual effects on brain activity in sensory cortices within the theoretical framework of predictive coding (PC framework), which emphasises the importance of prediction◻error coding and enhanced neural representations of unexpected stimuli^2–5^.

The resulting empirical landscape, however, remains difficult to reconcile into a coherent account of how expectations shape sensory processing^6–9^. Most notably, conflicting evidence suggests that expectations can lead to either enhanced or reduced fidelity of sensory representations of expected stimuli, processes referred to as ‘sharpening’ and ‘dampening,’ respectively^10–14^. This divergence leaves open a fundamental question: does expectation exert a ‘prediction-error’ effect by suppressing redundant information, or a ‘Bayesian’ effect by selectively enhancing the precision of expected signals?

At the neural level, sharpening models propose that expectations suppress neurons not tuned to the expected stimulus, thus inhibiting inconsistent information. This results in sharper tuning curves, increasing the signal-to-noise ratio for expected stimuli^15^. In contrast, 1) dampening accounts posit that expectations suppress neurons that are tuned to the expected stimuli^16^, and 2) prediction error accounts posit that expectations enhance neurons that are tuned to the unexpected stimuli^17^. Studies attempting to settle this debate have produced mixed results, largely due to a reliance on functional magnetic resonance imaging (fMRI). Some fMRI and multivariate pattern analysis (MVPA) studies report neural sharpening for expected visual information^11,13,18,19^, while others find evidence for neural dampening, indicated by lower stimulus decoding^12,20^ or rather prediction error, i.e., stronger unexpected compared to expected representations^13,17,21,22^.

To resolve this perceptual prediction paradox, the Opposing Process Theory (OPT) proposes that sharpening and dampening both occur, but at distinct time points^16^. Based on behavioral trends showing that expected events are perceived with greater intensity early on but reverse, favoring unexpected events later, OPT suggests that initial processing relies on prior knowledge to sharpen sensory representations, followed by a later reactive stage that dampens expected representations to highlight surprise. Enhanced early representations of expected stimuli are thought to be facilitated by a pre-activation of the corresponding sensory template^23–26^, although the conditions under which such pre-activation occurs remain contested^27,28^. Recent electrophysiological research has begun to leverage the high temporal resolution of EEG to parse out these dynamics and validate the OPT. For instance, McDermott and colleagues^29^ investigated a visual statistical learning task and found that within a trial, expectation increased decoding accuracy at early latencies (sharpening) but decreased it at later latencies (dampening). These effects fluctuated dynamically across trials, suggesting that sharpening and dampening may emerge at distinct stages of associative learning.

In the auditory domain, we recently demonstrated that sharpening and prediction error processes co-exist during speech comprehension, but operate at distinct levels of processing^14^. By integrating EEG speech decoding with transformer-based encoding models, we found that listeners’ speaker-specific semantic priors sharpened low-level sensory representations, pulling them toward expected acoustic signals. Conversely, prediction errors simultaneously emerged at higher linguistic levels. This hierarchical dissociation supported a unified model wherein lower-level sharpening facilitates robust perception while higher-level prediction errors support adaptive world models.

The present study seeks to further elucidate the hierarchical and temporal coordination of these opposing mechanisms in a highly controlled visual experiment that uses composite images composed of both an expected and a surprising component. This paradigm, in conjunction with time-resolved mass univariate linear discriminant analysis (LDA), allowed us to specifically test whether expectation facilitates an initial Bayesian effect followed by a subsequent prediction◻error signaling phase, consistent with the dual◻process theory proposed by Press and colleagues^16^. To this end, we measured how expectations influenced the extent to which EEG signals encode image-identity information over time, presenting four natural images (car, house, cat, elephant) in a predictable sequence interspersed with composite images that combined one expected and one unexpected image.

Hierarchical predictive processing accounts propose that the cortical hierarchy represents the visual world at multiple levels of abstraction simultaneously, with more superordinate information represented in higher-order areas ^30^. Several studies, however, extend this framework, showing that prediction error signals in lower levels predominantly reflect the deviance of higher-level stimulus features^17,31,32^. How expectations affect sub- vs. superordinate stimulus representations over time is still an open question. Although our paradigm was not optimized to answer this, we leveraged the superordinate distinction in our image set (two animal images vs. two images containing no animals) to obtain initial insights into how expectation differentially affects sub- and superordinate image representations over time.

## Results

During our EEG study, participants viewed four natural images depicting a car, a house, a cat, and an elephant. During each 1.5◻hour EEG session, these images were presented in a fixed repeating order (e.g., car–house–cat–elephant–car–house–cat–elephant…, as shown in Figure 1a). Participants viewed this sequence more than 2,500 times per session, rendering the occurrence of each image highly predictable. Each participant completed six EEG sessions, across which they were shown all six possible unique repeating image sequences. Data analysis was performed across all six sessions for each participant to rule out low◻level perceptual sequence effects, including adaptation◻related transition-patterns^8^. Participants were instructed to continuously fixate the central fixation cross and, to ensure that they remained vigilant and attended to all images, we asked them to press a button whenever an upside-down image was presented. These catch trials were shown during 1.6% of all trials, and participants detected them during 98.25% (SD = 1.69%) of all trials with an average reaction time of 663 ms (SD = 79 ms).

**Figure 1.**
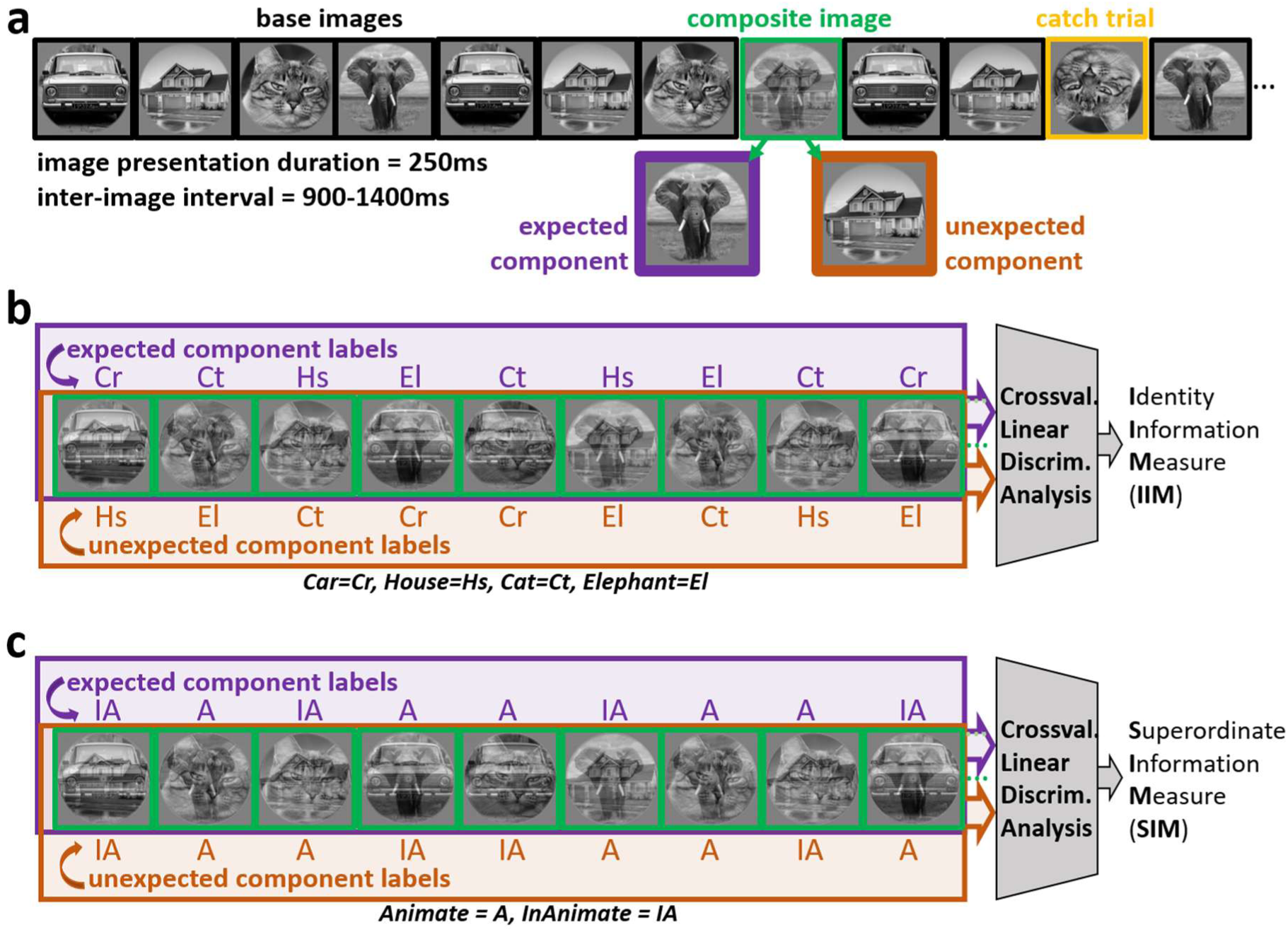
**Experimental design and data analysis *a EEG experimental paradigm.*** Participants viewed a continuous, highly predictable stream of four base images presented in a fixed repeating order (e.g., car→house→cat→elephant→car→house…). Each image was presented for 250 ms, separated by a variable inter-image interval of 900–1400 ms. In 24% of trials, a composite image (green outline) was presented, formed by superimposing the predicted image (expected component; purple outline) and a surprising image (unexpected component; brown outline). Participants maintained vigilance by detecting infrequent (1.6%) catch trials depicting upside-down images (yellow outline). ***b Identity Information Measure (IIM)*** To measure neural processing of image identity during composite image presentations, cross-validated Linear Discriminant Analysis (LDA) was applied to EEG data. Classifiers were trained and tested separately using labels corresponding to the identity of the expected component (purple upper box) or the identity of the unexpected component (brown lower box). ***c Superordinate Information Measure (SIM),*** analogous to the IIM analysis, LDA was used to quantify ***superordinate image*** processing, here defined by animacy. Composite images were labeled according to whether their expected components (purple upper box) or unexpected components (brown lower box) depicted an animal.

Critically, during 24% of image presentations, we showed a composite image consisting of the expected image and an unexpected image (Figure 1a). These composite images allowed us to examine image◻identity processing separately for the expected and unexpected components by labeling each composite according to either the identity of its expected or unexpected component (Figure 1b). Moreover, to examine the processing of superordinate image information, we also labeled the composite images according to whether their expected and unexpected components depicted an animal (Fig 1c). To measure these two types of image component information processing, we applied Linear Discriminant Analysis (LDA) separately for each electrode and labeling scheme to quantify each component’s Identity Information Measure (IIM) and Superordinate Information Measure (SIM) over time. These two measures were computed based on the LDA-based cross◻validated posterior probability values of image components, as described in more detail in the Star★Methods Section.

As a sanity check, we first quantified IIMs and SIMs for the base images. Consistent with previous work^33,34^, this analysis revealed persisting decodability of base◻image identity and superordinate image information, emerging at 56 ms and 132 ms, respectively, in clusters spanning all EEG electrodes (**IIM**: *p*FWE < .01; *t*_sum_ = 119904.1, *t*_range_ = 2.34 to 84.54; IIM_range_ = 0.0001, 0.0311; **SIM**: *p*FWE < .01; *t*_sum_ = 48230.8, *t*_range_ = 2.33 to 68.99; SIM_range_ = 0.0001–0.0399). These effects were also observed for the expected and unexpected image components (Figure 3 a,b,d,e), albeit overall less strongly and later than for the base trials, likely because the LDA classifier had to extract information from lower◻contrast images while simultaneously unmixing them from the overlapping component images.

In addition, given that this analysis is based on EEG responses to 10,200 highly predictable images presented to each participant, it offers a highly powered test for representations of expected stimuli before stimulus onset. However, it revealed no evidence for pre-stimulus representations of either identity or superordinate information about the upcoming image. To determine whether this null-finding was due to baseline correction subtracting out pre-stimulus effects, or due to the statistical threshold being too conservative as a result of a large number of multiple comparisons, we repeated it for non-baseline corrected data specifically for the timepoints between -300 and 0 ms and using a more liberal cluster forming and FWE threshold of α = .05. With this analysis, we also did not find any significant identity or superordinate information before image onset (**max IIM cluster:** *p*FWE = .527, *t*_sum_ = 70.14; **max SIM cluster:** *p*FWE = .830, *t*_sum_ = 28.82).

Next, we turned to the composite images. For the expected image components (Figure 3a), identity information rose significantly above chance between 92 and 700 ms in a widespread cluster of 53 electrodes (*p*FWE < .01; *t*_sum_ = 18902.7, *t*_range_ = 2.33 to 14.70; IIM_range_ = 0.0002 to 0.0041). For the unexpected components (Figure 3b), identity information exceeded chance between 108 and 700 ms in a widespread cluster of 59 electrodes (*p*FWE < .01; *t*_sum_ = 21654.4, *t*_range_ = 2.33 to 13.73; IIM_range_ = 0.0002 to 0.0037). A direct comparison of IIM values for expected versus unexpected components (Figure 3c) revealed two clusters of expectation◻related differences. The earlier cluster of 47 electrodes emerged between 288 and 380 ms, centered on the right parieto◻occipital region, showing enhanced processing of expected component identity (*p*FWE < .01; *t*_sum_ = -1174.6, *t*_range_ = -2.58 to -6.93; IIM Difference_range_ = -0.0003 to -0.0024). In contrast, the later cluster (424–700 ms) showed enhanced processing of unexpected component identity across a cluster of 44 electrodes, centered on the left parieto-occipital and frontal region (*p*FWE < .01; *t*_sum_ = 1892.1, *t*_range_ = 2.58 to 6.12; IIM Difference_range_ = 0.0002 to 0.0013). An individual analysis of single-subject effects revealed that within both the earlier and the later cluster, all six participants displayed the reported effect, while a Dixon’s Q test did not reveal evidence for either effect being driven by an outlier (earlier cluster: Qmax = 0.435, p > .05; later cluster: Qmax = 0.346, p > .05; Supplemental Figure 1). Together, these findings indicate that expectation initially boosts identity processing for expected components, followed by enhanced processing of unexpected component identity at later stages. Having established these expectation effects on component identity representations, we next examined whether similar effects emerge for representations of more superordinate information by assessing the processing of animate vs. inanimate image components.

For expected components (Figure 3d), superordinate image information rose significantly above chance level in three clusters occurring during the time windows 180–244 ms, 288–404 ms 412–548 ms (all *p*FWE < .01; **cluster 1**(30 electrodes): *t*_sum_ = 593.1, *t*_range_ = 2.33 to 6.10; SIM_range_ = 0.0003 to 0.0020; **cluster 2**(35 electrodes): *t*_sum_ = 1706.3, *t*_range_ = 2.33 to 7.98; SIM_range_ = 0.0003 to 0.0033; **cluster 3**(20 electrodes): *t*_sum_ = 946.4, *t*_range_ = 2.33 to 5.15; SIM_range_ = 0.0003 to 0.0015). For unexpected components (Figure 3e), superordinate image information rose significantly above chance level in three clusters occurring during the time windows 176–236 ms, 340–448 ms, and 448–520 ms (all *p*FWE < .01; **cluster 1**(36 electrodes): *t*_sum_ = 595.8, *t*_range_ = 2.35 to 7.98; SIM_range_ = 0.0003 to 0.0033; **cluster 2**(40 electrodes): *t*_sum_ = 2113.2, *t*_range_ = 2.33 to 8.42; SIM_range_ = 0.0003 to 0.0032; **cluster 3**(34 electrodes): *t*_sum_ = 646.2, *t*_range_ = 2.33 to 4.84; SIM_range_ = 0.0003 to 0.0014). A direct comparison of SIM values (Figure 3f) revealed two clusters of expectation◻related differences. The earlier cluster of 28 electrodes emerged between 288 and 352 ms, centered on the right parieto◻occipital region, showing enhanced processing of expected superordinate information (*p*FWE < .01; *t*_sum_ = -826.1, *t*_range_ = -2.59 to -6.76; SIM Difference_range_ = - 0.0004 to -0.0035). In contrast, the later cluster (356–420 ms) showed enhanced processing of unexpected superordinate information across a cluster of 32 electrodes, centered on the left temporo-parietal region (pFWE < .01; *t*_sum_ = 588.4, *t*_range_ = 2.58 to 5.98; SIM Difference_range_ = 0.0005 to 0.0025). An individual analysis of single-subject effects revealed that within the earlier cluster, five of six participants displayed the reported effect, while all participants displayed the effect reported for the later cluster. Again, we found no evidence for the presence of outliers (earlier cluster: Qmax = 0.279, p > .05; later cluster: Qmax = 0.527, p > .05). These findings indicate that, as for identity information, expectation also initially boosts superordinate information for expected components, followed by enhanced unexpected superordinate information at a later stage.

Exploring whether sub- and superordinate representations are affected differently over time by expectation revealed a cluster of 29 electrodes, centered on the left parieto-occipital region between 448 and 544ms (Figure 3g), in which expectation effects were greater for identity than for superordinate information (*p*FWE < .01; *t*_sum_ = -556.0, *t*_range_ = -2.58 to -6.02; Z-score-modulation_range_ = -0.0245 to -0.1036). In addition, there was a trend (Supplemental Figure 2) for greater modulation of superordinate vs. subordinate information during an earlier time window between 356 and 416ms (*p*FWE = .014; *t*_sum_ = 426.3, *t*_range_ = 2.59 to 4.96; Z-score-modulation_range_ = 0.0204 to 0.0796). Taken together, these findings suggest that expectation violations give rise to an earlier enhancement of representations of superordinate as compared to subordinate image properties.

Finally, to compare our findings with previously reported effects of expectation on EEG responses, we conducted two additional analyses. First, we explored whether expectation effects reverse across trials^29^. To this end, we tested whether expectation effects on identity and superordinate information, within the clusters shown in Figure 3c and Fig 3f, differ across the five 11-minute runs within each EEG session. However, none of the four fixed-effects ANOVAs testing for an effect of run revealed a significant effect (F(4,175) < 0.728, all p > .58; effect sizes across runs, clusters, and information measures are shown in Supplemental Figure 3). Therefore, our data provide no additional evidence that expectation effects change across trials. Second, though recent EEG studies investigated effects of expectation with decoding based on response patterns across all EEG sensors^29,35^, an LDA analysis trained on patterns across all electrodes did not reveal any significant clusters for expectation effects in our data (**max IIM cluster**: *t*_sum_ = 33.24, *p*FWE = .087; **max SIM cluster**: *t*_sum_ = 7.67, *p*FWE = .607, Supplemental Figure 4). This suggests that multivariate analyses across all EEG electrodes can miss robust effects within single electrodes, which may help explain why other EEG studies found no effect of expectation^35^.

## Discussion

Taken together, our results reveal a consistent temporal pattern of how expectations affect representations of image identity and superordinate image information over time. The processing of both types of information exhibits an initial enhanced representation of the expected image components, suggesting an earlier phase during which prior knowledge is integrated with sensory input to support veridical perception. This earlier facilitation was followed by a later phase during which processing of both types of information was stronger for the unexpected components, indicating increased processing of surprising information. This biphasic pattern, therefore, supports the proposal that perceptual expectations first promote Bayesian integration to optimize current perceptual inferences, and subsequently amplify unexpected events to refine future predictions^16^. Thereby, our findings contribute to the ongoing debate on how expectations shape sensory processing^6–9^.

When comparing the temporal dynamics of expectation effects on identity and superordinate image information, a difference becomes apparent. Although expectation enhanced the processing of expected component identity and superordinate information during a comparable temporal window, the subsequent enhancement for unexpected components appeared to unfold differently. Specifically, enhanced superordinate information for unexpected components emerged earlier than identity information, as partially confirmed by our direct analysis of the difference between how expectation affects identity vs. superordinate information processing. These findings suggest that surprising stimuli first trigger an increased awareness of the kind of unexpected event that occurred, e.g., its broad, categorical “gist”, before the system resolves the more fine◻grained perceptual details needed to identify what exactly was surprising. Therefore, our results are consistent with the notion that prediction error signals, evoked during earlier sensory information processing, are best explained by the deviance of superordinate stimulus features^31,32^. In a broader sense, our findings and these previous findings resonate with the “forest before the trees” principle articulated in reverse hierarchy theory^36,37^, in which coarse, global representations are accessed earlier and guide the subsequent refinement of detailed perceptual interpretations.

All expectation◻related effects we observed emerged no earlier than 288 ms after stimulus onset, far too late to reflect modulation of the initial feedforward sweep. This timing is later than recently reported expectation effects on EEG responses^29,38^, but it aligns with studies suggesting that expectation effects are predominantly cognitive rather than strictly perceptual^7,39^. Moreover, our findings align with recent proposals that prediction errors primarily reflect updates based on higher-level information across the processing hierarchy, rather than from low-level sensory discrepancies^17,32,40^.

The temporal profile of our effects also overlaps with the highly discussed N400 component^41^. Although interpretations of the N400 vary, from semantic retrieval and contextual integration to signaling prediction error, contemporary accounts increasingly view it as a graded index of contextual facilitation and prediction◻confirmation during semantic access^42,43^. The succession of opposing expectation effects we observe is consistent with these perspectives: expectations modulate brain activity through a biphasic mechanism, with an initial enhancement of processing of expected image components followed by a later prioritization of unexpected components. This temporal pattern suggests a dynamic interplay between facilitation driven by prior knowledge and subsequent prediction◻error computations at higher categorical levels, consistent with Bayesian accounts of perception and enabled by recurrent interactions within the visual hierarchy^44^. Our findings also coincide temporally with recurrent high◻level effects in the lateral occipital cortex, indicating that prior◻induced top◻down processing can enhance representations of expected stimulus features^45^.

By analyzing EEG data during a statistical learning task, a recent EEG study found that expectation enhances neural processing early in a single trial and suppresses it later, but exhibits the reverse pattern across the overall learning process^29^. Specifically, over the course of the experiment, expectation initially decreased neural decoding accuracy during earlier learning trials and then increased it in later trials. These results suggested that opposing mechanisms operate hierarchically at distinct stages of associative learning. Our study, however, did not reveal any evidence for expectation effects increasing or decreasing across the five 11-minute runs of each EEG session. These conflicting findings, therefore, indicate that future studies are needed to reveal how the acquisition of knowledge over time translates into the Bayesian and prediction-error effects.

The temporal sequence observed in our data, an earlier facilitation of expected information followed by a later enhancement of unexpected information, is compatible with models of serial template matching^46^. Under this framework, the visual system first aligns incoming sensory input with a pre-activated template of the anticipated image, while subsequently redirecting resources toward processing the unpredicted aspects of the image. The ability to successfully apply a specific template upon stimulus onset inherently requires the observer to have implicitly learned the sequence statistics to generate a prior. Though previous research suggests that prior expectations induce sensory templates^23,24,47^, it remains a matter of debate whether such preactivations exist^27,28^, and our study also does not provide evidence for the existence of sensory templates emerging before, or while, predictable images are presented, despite the high-powered analysis we used to assess this, which was based on more than ten thousand trials per participant. Therefore, further research is needed to provide a conclusive understanding of whether, and if so, under which conditions, i.e., under which tasks and behavioural relevance^48^ or explicit versus implicit priors, pre-activated sensory templates emerge.

### Limitations

A primary limitation of the current study is the small sample size (N = 6), which notably includes all three of the study’s authors. This introduces potential sampling bias and limits the generalizability of the findings. However, this constraint is mitigated by the depth of the data collection. By testing this small cohort six separate times, we were able to generate a highly balanced within-participant dataset that maximizes internal consistency. In addition, the studied context effect based on four repeating images is unlikely to be subject to voluntary influence or intentional bias. Accordingly, three of the four main effects of expectation reported in this study were present in all six participants, with the fourth effect being present in five out of six participants (Supplemental Figure 1).

An additional limitation is the 24% frequency of composite trials. While necessary to yield enough data for our analysis, this relatively high proportion means participants may have begun learning the composite images, which could weaken the ‘highly predictable’ structure of the sequence or render the composites predictable themselves. Importantly, such learning would work against our hypotheses by reducing the difference between unexpected and expected image components.

Another limitation of the present study is the inherent difficulty in disentangling content-specific prediction errors from generic, deviance-driven attentional reorientation. The design of our study minimizes attention confounds by intentionally omitting a task associated with the expected and unexpected components. This choice reflects a deliberate methodological compromise: by prioritizing clean, unconfounded neural signatures of expectation, we accepted the trade-off of refraining from behavioral assessments of the expectation effects. Nonetheless, the transition probabilities, which defined the expected and unexpected conditions, remained stable within each run. Because this environment did not require active model updating, the late enhancement observed for the unexpected image component might not purely reflect a model-updating prediction error. Instead, it could be driven by an attentional shift toward a surprising or unfamiliar visual element^49,50^. Separating true predictive surprise from bottom-up attentional capture is a well-recognized and persistent challenge in expectation research, as violations of expectations naturally recruit attention^48^. Nevertheless, whether this late processing stage is primarily driven by model-updating prediction errors or by surprise-induced attentional reorientation, our core conclusion remains unaffected: the visual system exhibits a distinct temporal segregation, prioritizing expected and unexpected visual information sequentially.

Our stimulus set was restricted to four images, with only two exemplars per superordinate category (animate, inanimate). This limited sampling constrains the generalizability of our conclusions regarding superordinate versus identity-level processing, as category-level effects estimated from only two exemplars may be confounded with idiosyncratic, image-specific features rather than reflecting genuine animacy-based representations. A larger stimulus set, with multiple exemplars per category, would be required to disentangle category-general effects from stimulus-specific ones and to more firmly establish the hierarchical timing differences we observed between superordinate and identity representations.

Finally, we would like to acknowledge a potential effect of stimulus-specific adaptation. Although we mitigated adaptation by ensuring that component images were never presented in the preceding trial (i.e., 1-back), the presentation history was not completely equated for longer lags. Specifically, the expected component image was consistently presented four items before, whereas the unexpected item appeared either two or three items before. Consequently, the unexpected component may have been subject to stronger neural adaptation, a well-documented confound with expectation effects^51^. This effect, however, cannot explain our finding of a later enhancement of unexpected-component processing, as stronger adaptation for unexpected components should reduce, and not enhance, the relative strength for unexpected components. In addition, the enhanced processing of identity and superordinate information for expected components reported in this study appears to be unlikely to be fully accounted for by an adaptation effect, as adaptation has been shown to have its maximal effect on visually evoked EEG responses between 130ms and 180ms after stimulus onset^52^, while our effects emerge no earlier than 288ms after image onset.

## Conclusion

This EEG study reveals that expectations shape visual processing through a biphasic mechanism: the brain first uses prior knowledge to enhance the processing of expected information, and then shifts to amplify surprising events. Therefore, our data suggest that Bayesian integration of sensory information and enhanced processing of surprising events unfold sequentially rather than simultaneously in the human brain. For unexpected stimuli, our results provide additional support for the notion that the brain prioritizes the processing of coarser superordinate image information before resolving their detailed identity, a pattern that echoes the principles of reverse◻hierarchy theory, in which coarse, global representations precede fine◻grained perceptual detail. Finally, the relatively late timing of the expectation effects observed in this study supports recent proposals that expectation effects are more ‘cognitive’ than ‘perceptual’ in nature^7^.

## Resource availability

## Lead Contact

Further information and requests for resources should be directed to and will be fulfilled by the lead contact, Arjen Alink

## Materials Availability

This study did not generate new unique reagents.

## Data and Code Availability

- Raw EEG data, MATLAB preprocessing, analysis and plotting scripts, stimulus files, and the Python script used for stimulus presentation have been deposited at the Open Science Framework (OSF) and are publicly available as of the date of publication. The DOI/link is listed in the Key Resources Table.
- Any additional information required to reanalyze the data reported in this paper is available from the lead contact upon request.

## Acknowledgments

We thank Antoniya Boyanova, Ivana Tanasic, and Nahid Hasan for constructive feedback regarding the manuscript. We acknowledge support by the European Research Council (ERC2021-CoG-101044616 to AA). This work is funded by the Emmy Noether program of the Deutsche Forschungsgemeinschaft (German Research Foundation; Grant No DFG BL 1736/1-1 to HB). This work was supported by the Research Network for the Interdisciplinary Study of Predictive Processing, funded by the Deutsche Forschungsgemeinschaft (DFG) - Project number 442810744.

## Author contributions

Conceptualization, A.A. and H.B.; methodology, A.A.; investigation, A.A., J.B., and H.B.; visualization, A.A.; funding acquisition, A.A and H.B.; project administration, A.A. and H.B.; supervision, A.A and H.B.; writing, A.A and H.B.;

## Declaration of generative AI and AI-assisted technologies in the writing process

During the preparation of this work, the authors used ChatGPT-4.0 and Gemini 3 to improve style and readability. After using this tool/service, the authors reviewed and edited the content as needed and take full responsibility for the content of the published article.

## STAR★Methods

## Key resources table

| Reagent or Resource | Source | Identifier |
| --- | --- | --- |
| **Deposited Data** |  |  |
| Raw EEG data, stimulus files, and analysis scripts | This paper; OSF | https://osf.io/bdtfr/overview?view_only=e7683c7602f7470e83a5e0683ed98223 |
| **Software and Algorithms** |  |  |
| MATLAB Release R2021b | The MathWorks, Inc. | RRID: SCR_001622 |
| EEGLAB toolbox | Delorme & Makeig (2004) | RRID: SCR_007292 |
| PsychoPy | Peirce et al. (2019) | RRID: SCR_006571 |
| SHINE toolbox for MATLAB | Willenbockel et al. (2010) | https://github.com/mwillenbockel/SHINE |
| FieldTrip | Oostenveld et al., 2011 | RRID:SCR_004849. |
| BrainVision Recorder | Brain Products | N/A |
| **Other** |  |  |
| 64-channel ActiCap | Brain Products, Gilching, Germany | N/A |
| 23-inch TFT-Screen (SyncMaster P2370) | Samsung, Seoul, South Korea | N/A |

## Experimental model and study participant details

### Human Subjects

Six participants took part in six EEG sessions (three females and three males, age range 19 to 38 years) between 18.02.2021 and 21.08.2021. All three authors participated in this study. The research was approved by the local ethics committee (Ethics Committee of the Chamber of Physicians in Hamburg, approval no. PV7210, approval date: 18.02.2020), and all participants provided written informed consent.

## Method details

### Stimulus Generation and Presentation

Stimuli consisted of four greyscale natural images of a cat, an elephant, a car, and a house. Images were downloaded from the Pixabay and Unsplash stock photo websites under their free-to-use licences. Images were transformed into grey-scale images, and we applied a fade-out mask to them to make the images’ peripheral and central areas grey. Subsequently, we reduced low-level differences across images by applying the spectrum, histogram, and intensity normalisation and equalisation (SHINE) toolbox^53^ for MATLAB. All images were presented using PsychoPy on a 23-inch TFT-Screen (SyncMaster P2370; Samsung, Seoul, South Korea) positioned centrally 110 cm in front of the participant. The circular stimulus area had a radius of 7.24-degree visual angle (°), the central grey area had a radius of 0.59°, and the black fixation dot had a radius of 0.08°. Composite images were created while running PsychoPy by drawing one image in the background and drawing the second image with 50% opacity on top of it. All base images were presented with 75% opacity.

We presented these stimuli to participants in a deterministic, fixed repeating sequence (as shown in Figure 1) during six 1.5-hour EEG sessions on six different days. Each of these sessions was subdivided into five ten-minute runs to accommodate short breaks for our participants. We presented each of the six possible repeating sequence orders (1234, 1243, 1324, 1342, 1423, and 1432) once to each participant (n = 6) during one of the six EEG sessions. Note that one can present a fixed repeating sequence of four images in six and not in twenty-four ways because, e.g., the image orders 1234, 2341, 3412, and 4123 give rise to the same repetitive sequence (…1234123412341234…). As a result, each of the four base images and each image component had a perfectly balanced stimulus history across all six sessions. The first image within a sequence was pseudorandomly selected such that each of the four images was shown at least once, but no more than twice, as the first image of a sequence across the six sessions. During a trial, an image was presented for 250 ms, followed by a 900-1400 ms inter-stimulus interval, during which only the grey background and the black fixation dot were presented. Participants were instructed to fixate the central fixation dot throughout the experiment and to press a button whenever a catch trial (upside-down base image) was presented.

Note that twelve unique composite images entail the six possible image pairs presented under two different expectation conditions. Hence, the composite image containing the elephant and the house, shown in Figure 1, was presented when a house was expected and when the elephant was expected. Composite images were pseudo-randomly interspersed in the image sequences while ensuring that each composite image was preceded by at least two base images. To preclude the effects of direct repetition, neither the expected nor the unexpected component of each composite image was based on the preceding base image. As a result, across all six sessions, physically identical composite images had the same balanced one-back history across expectation conditions. E.g., cat-house composites were equally often preceded by a car and an elephant stimulus, both for composite images presented when a cat and when a house image was expected.

## EEG Recording and Preprocessing

EEG data were sampled at 250 Hz using 64 active electrodes positioned according to the extended 10–20 system (ActiCap, Brain Products, Gilching, Germany) and recorded with BrainVision Recorder. Preprocessing was performed in MATLAB (The MathWorks, Inc. MATLAB Release R2021b. Natick, Massachusetts, USA (2021), using the EEGLAB toolbox^54^, and included standard high◻ and low◻pass filtering at 0.1 Hz and 100 Hz, respectively. The continuous data were then epoched (-200 to 800 ms) and baseline◻corrected using the average signal from −200 ms to stimulus onset. No additional preprocessing or artifact◻rejection procedures were applied, as such steps are more likely to reduce than enhance statistical power^55^.

## Quantification and statistical analysis

Because expectation effects in EEG can be subtle and difficult to detect^35^, we collected a large amount of data per participant: 2,400 composite◻image trials and 10,200 base◻image trials for each of our six participants. Data were analysed using a fixed◻effects approach, providing high statistical power while acknowledging that the effects cannot be expected to generalize beyond the measured sample.

We performed the LDA analysis separately for each participant, using 10◻fold cross◻validation, to obtain posterior probability values for left-out base images and for left-out expected and unexpected components of the composite images. Posterior probabilities for composite images were computed after labeling them as either their expected or unexpected components (Figure 1), yielding image identity measures (IIMs). Analogously, labeling composite images by the animacy of their expected and unexpected components (cat, elephant vs. house, car) yielded raw superordinate image measures (SIM_raw). To obtain SIM values that more specifically reflect superordinate rather than idiosyncratic low◻level image features, we also computed SIM_control values based on the two alternative pairings (cat, house vs. elephant, car; cat, car vs. elephant, house). These SIM_control values contain no animacy information but capture the low◻level visual structure we aimed to control for. Final SIM values were therefore computed by subtracting the average SIM_control value from SIM_raw. All IIM and SIM values were computed in a mass◻univariate manner, separately for each timepoint and each electrode (excluding eye electrodes, n = 60).

During six of our eight analyses (Figure 2a,b, Figure 3a,b,d,e), we performed one-sided t-tests (positive only) to determine if observed IIM and SIM values were significantly greater than chance level and zero, respectively. For base images, this test was applied across all single◻image IIMs from all participants. For the expected and unexpected components of the composite images, two separate t◻tests were applied across all expected and unexpected component IIMs from all participants. For the remaining three analyses (Figure 3c,f,g), two-sided t-tests were performed versus zero. During two of these analyses (Figure 3c,f), we examined how expectation modulated IIM and SIM values and tested whether IIM- and SIM◻difference values (unexpected minus expected) were significantly greater or smaller than zero. During the third analysis (Figure 3g), we z-transformed observed IIM and SIM expectation modulations (across all trials, participants, timepoints, and electrodes within each information measure) and tested if the difference between z-transformed expectation modulations (SIM modulation minus IIM modulation) was significantly greater or smaller than zero for each electrode and timepoint.

**Figure 2.**
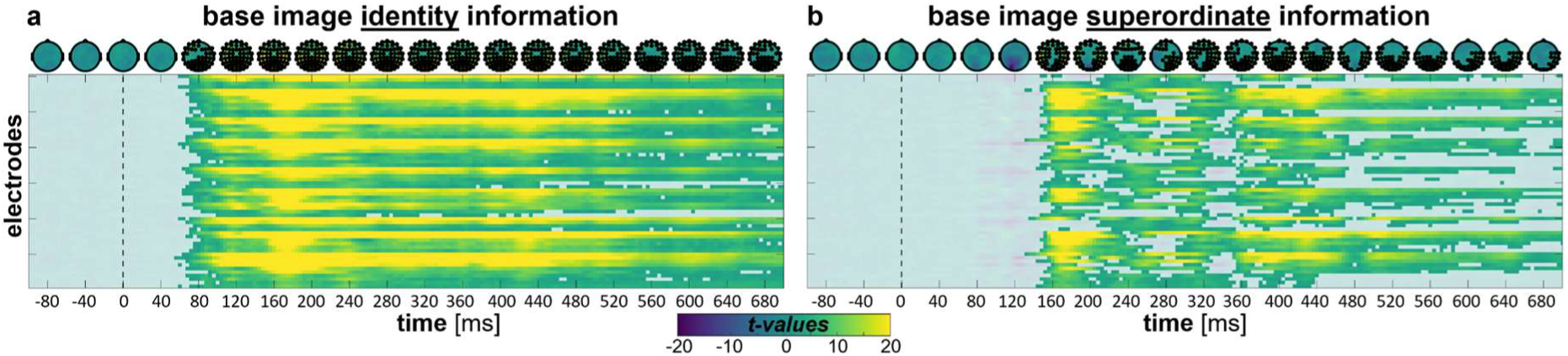
Identity and superordinate information of base images. Each matrix displays t-values, with rows representing individual electrodes and columns representing timepoints. Non◻significant t-values (*p*FWE > .01) are faded. Above each matrix, the same t-values are shown as topographic plots for selected timepoints from −80 to 680 ms in 40◻ms steps. Significant electrode locations in these topographic (*p*FWE < .01) plots are marked with a black dot. Matrix ***a*** shows t-statistics based on comparing observed base image identity measures (IIMs) to chance. The three electrodes with the highest t-values in the significant cluster (56–700ms) shown in this matrix are P8, PO8, and P6. Matrix ***b*** shows t-statistics based on comparing observed base image superordinate information measures (SIMs) versus zero. The three electrodes with the highest t-values in the significant cluster (132–700ms) shown in this matrix are P8, TP8, and P6.

**Figure 3.**
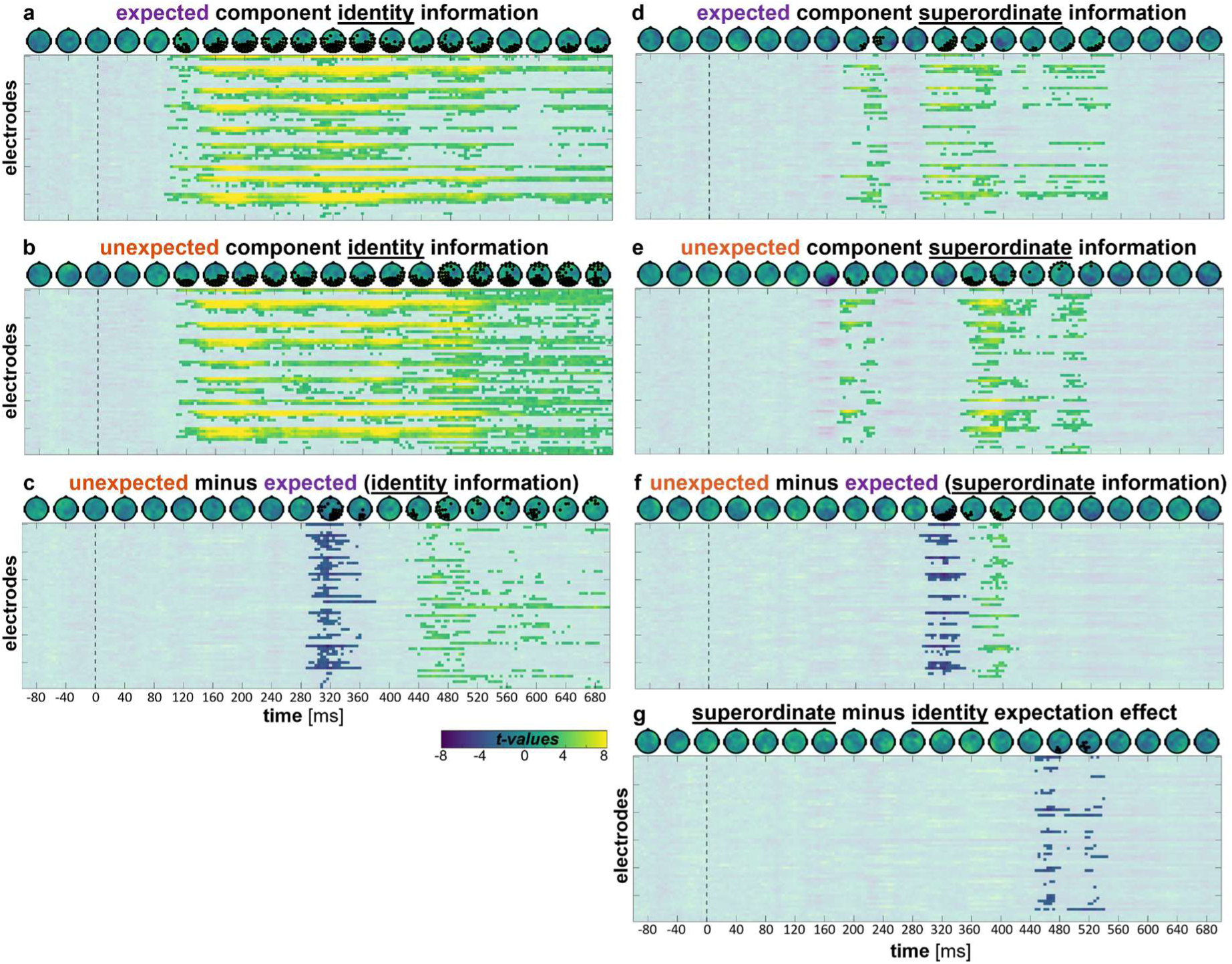
Component identity and superordinate information and their modulation by expectation. Each matrix displays t-values, with rows representing individual electrodes and columns representing timepoints. Non◻significant t-values (*p*FWE > .01) are faded in all plots. Above each matrix, the same t-values are shown as topographic plots for selected timepoints from −80 to 680 ms in 40◻ms steps. Significant electrode locations (*p*FWE < .01) in these topographic plots (exception, see g) are marked with a black dot. Matrices ***a*** and ***b*** show t-statistics based on comparing observed image identity measures (IIMs) to the chance level for the expected and unexpected components, respectively. Matrices ***d*** and ***e*** show t-statistics based on comparing observed image superordinate measures (SIMs) to zero for the expected and unexpected components, respectively. Matrices ***c*** and ***f*** display t-statistics for IIM and SIM difference values (unexpected minus expected). ***g*** displays the comparison of effects on identity vs. superordinate information. This test is performed across effect size differences (superordinate minus identity), computed after z-transforming within each information measure. The three electrodes with the highest t-values for all highlighted clusters are: **a:** (92– 700ms) P8, PO8, and P6, **b:** (108–700ms) P8, PO8, and P6, **c:** (288–380ms) P8, P6, and P4; (424–700ms) F2, PO3, and POz, **d:** (180–244ms) P6, PO8 and P8, (288–404ms) P6, PO4 and PO8, (412–584ms) Oz, PO8 and P6, **e:** (176–236ms) P8, P6 and TP8, (340–448ms) O1, Oz and PO7, (448–520ms) TP7, F3 and F5, **f** (288–352ms) P6, PO4, and P8, (356–420ms) P7, CP5, and P1, **g:** (448–544ms) PO3, POz and Oz.

To evaluate whether the magnitude of the neural expectation effects changed over the course of the EEG sessions, we analyzed the expectation effect sizes across the five 11-minute runs within each session for every significant cluster. To this end, we extracted the single-trial expectation effects for both the IIM and SIM information metrics. These trial-level data were then averaged within each run, yielding a mean effect size per run for each participant across their six respective sessions. To statistically assess run-related differences in expectation effect magnitude, we submitted these run-averaged effect sizes to a one-way fixed-effects analysis of variance (ANOVA), with chronological run number (Runs 1 through 5) designated as the sole fixed factor.

To correct for multiple comparisons across our spatiotemporal dimensions (200 timepoints and 60 electrodes), we applied a non◻parametric cluster◻mass permutation test (^56^, 1000 permutations; cluster◻forming threshold p < .01), controlling the family◻wise error rate (FWER) at a final significance level of p < .01. Cluster-mass was computed as the sum of t-values. Spatial clusters were defined based on electrode◻neighborhood relationships obtained via Delaunay triangulation of the 3D channel coordinates in FieldTrip (^57^; ft_prepare_neighbours, version 20260518), yielding a symmetric binary adjacency matrix that constrained cluster formation. To verify that our null findings regarding pre◻stimulus representations of information about the upcoming image were not due to overly conservative statistical thresholds, we repeated these analyses using a more liberal cluster◻forming and FWER threshold of p < .05.

## Supplemental information

**Supplemental Figure 1.**
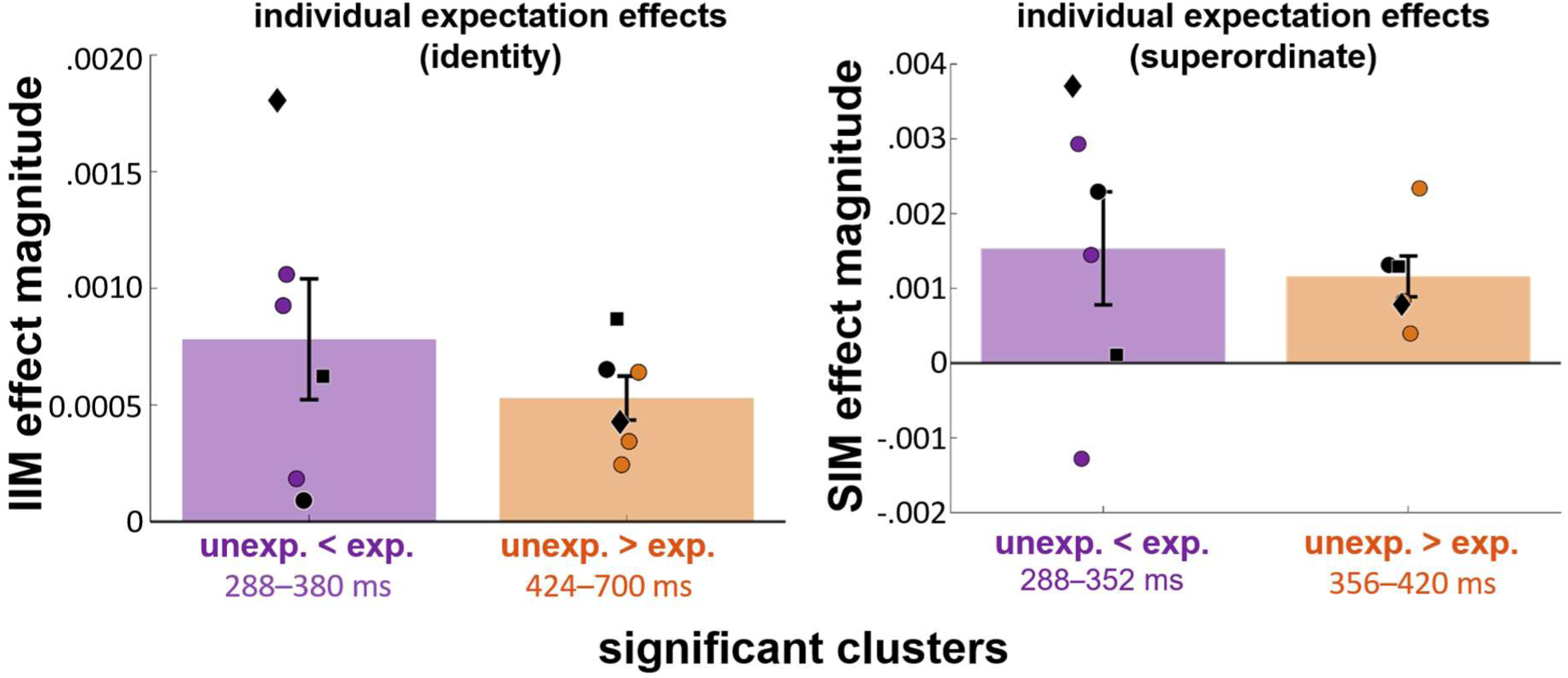
Expectation effects across participants. Bars and their SEMs reflect the effects of expectation across all participants within each of the four reported significant clusters shown in Figure 3c and Figure 3f. In addition, the average effect size for these clusters is shown separately for each of our six participants. All three authors were participants and are depicted, each with a unique black symbol (a black diamond, square, and circle). The other colored dots depict the three external participants.

**Supplemental Figure 2.**
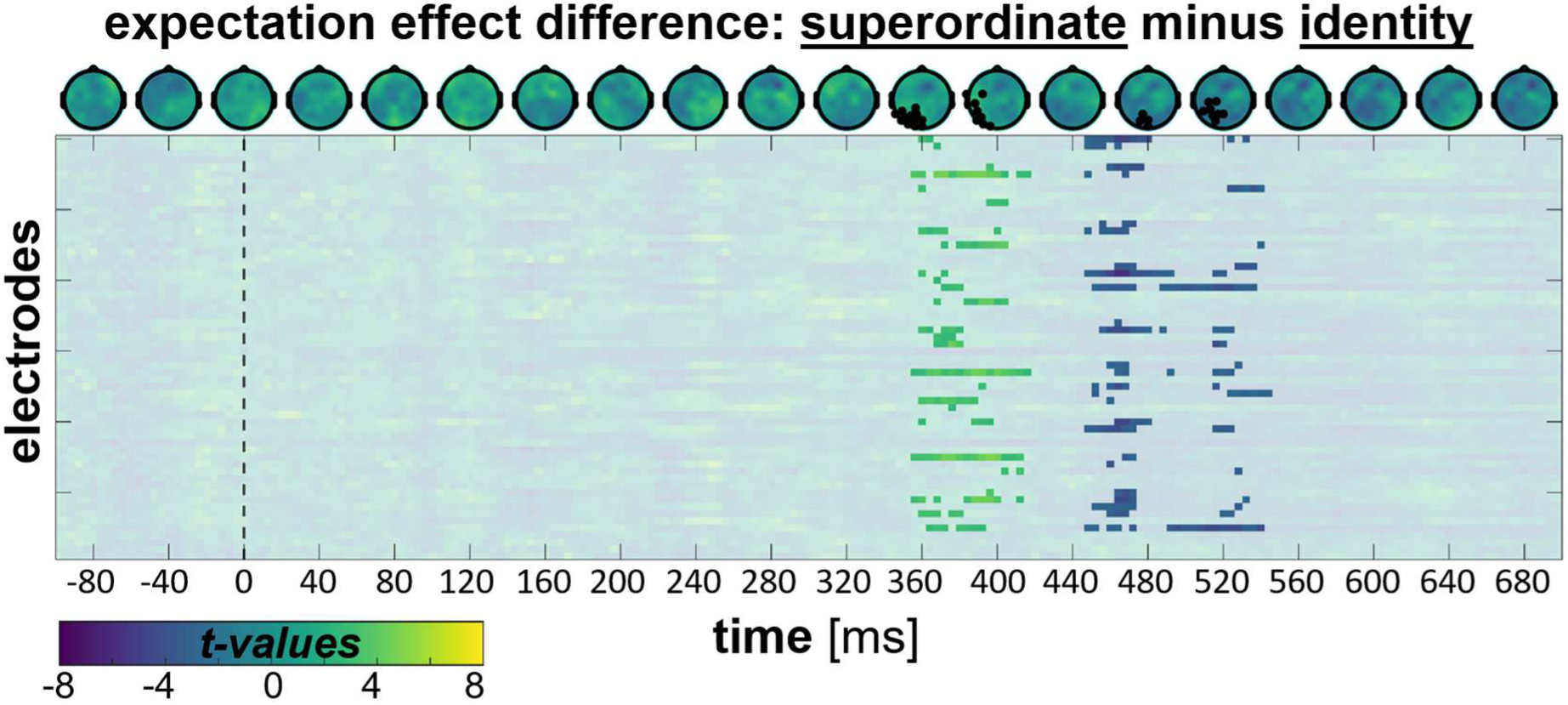
This figure displays results for the comparison of expectation effects on identity vs. superordinate information processing for the image components, similar to Figure 3g. These results, however, are based on a slightly less conservative threshold (*p*FWE < .02 instead of *p*FWE < .01). The three electrodes with the highest t-values for the two highlighted clusters are: (356–416ms) P7, PO7, and O1; (448–544ms) PO3, POz, and Oz.

**Supplemental Figure 3.**
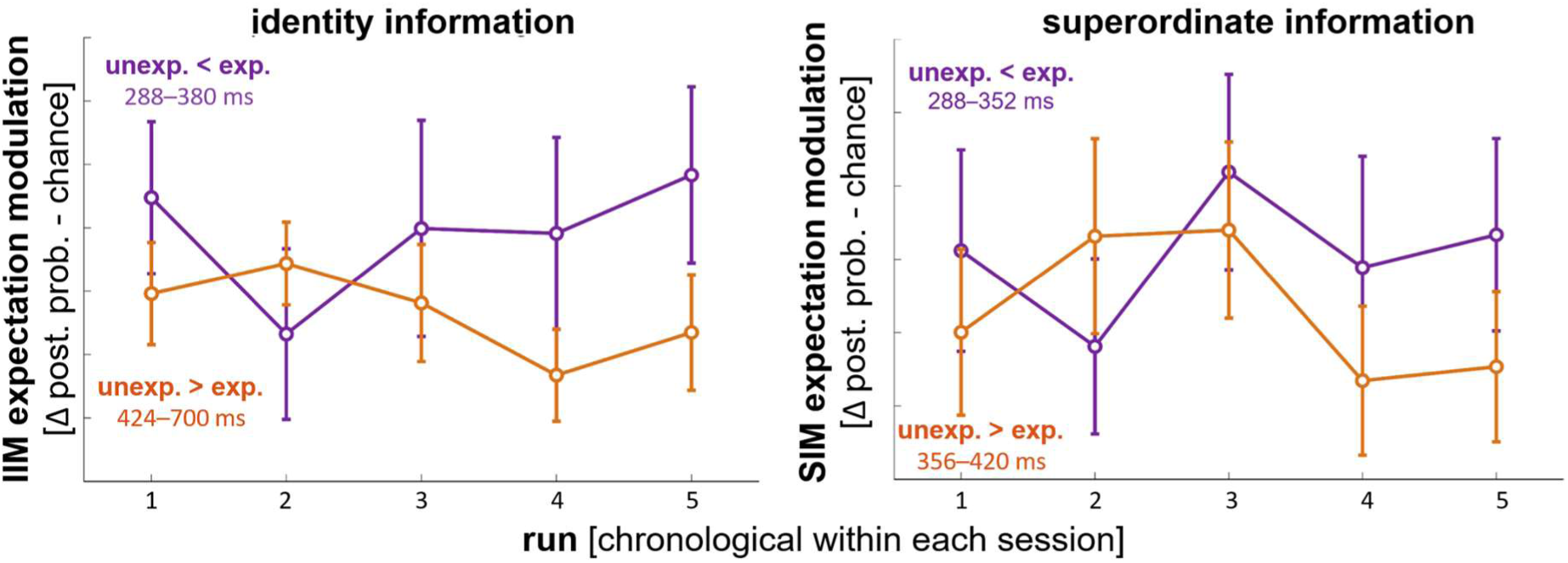
Temporal evolution of expectation effects across runs. Average effect size and its SEM, for each 11-minute run, for each of the four clusters shown in Figure 3c and Figure 3f. Averages and SEMs are computed across the six participants and the six EEG sessions, after averaging all trials within each run.

**Supplemental Figure 4.**
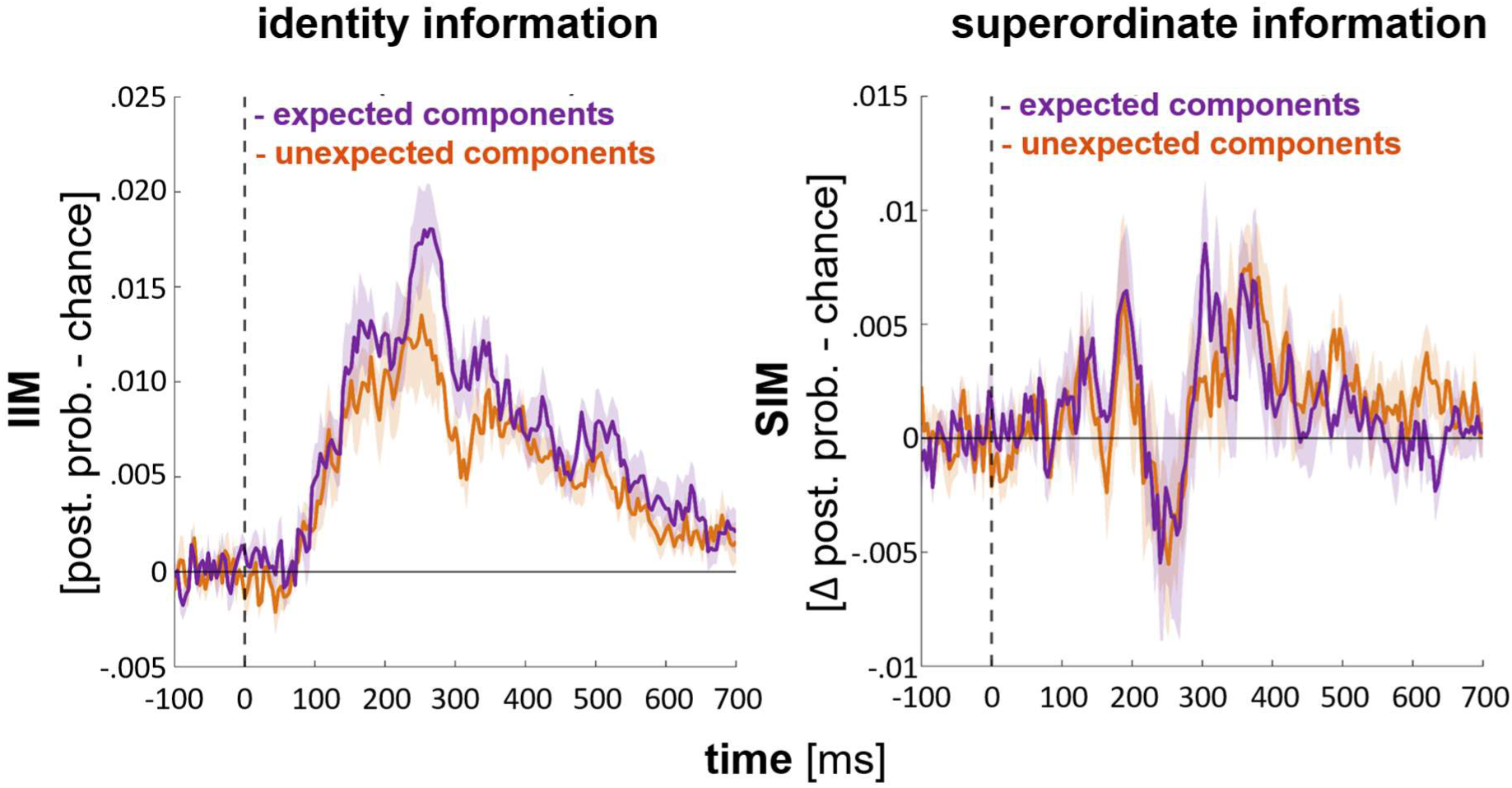
Component information measures based on LDA training and testing across all electrodes. Identity and superordinate information measures (IIM and SIM) time courses for expected and unexpected components based on LDA training and testing using EEG response patterns across all electrodes. In contrast to our mass-univariate single-electrode LDA analysis, this analysis does not reveal any statistically significant effect of expectation.

